# Depth and turbidity affect *in situ* pumping activity of the Mediterranean sponge *Chondrosia reniformis* (Nardo, 1847)

**DOI:** 10.1101/2020.03.30.009290

**Authors:** Mert Gokalp, Holger Kuehnhold, Jasper M. de Goeij, Ronald Osinga

## Abstract

Effects of depth and turbidity on the *in situ* pumping activity of the Mediterranean sponge *Chondrosia reniformis* (Nardo, 1847) were characterized by measuring osculum diameter, oscular outflow velocity, osculum density per sponge and sponge surface area at different locations around the Bodrum peninsula (Turkey). Outflow velocity was measured using a new method based on video analysis of neutrally buoyant particles moving in the exhalant stream of sponge oscula, which yielded results that were in good comparison to other studies. Using the new method, it was shown that for *C. reniformis*, oscular outflow had a location-dependent, in most cases positive relationship with oscular size: bigger oscules process more water per cm^2^ of osculum surface. Turbidity and depth both affected sponge pumping in a negative way, but for the locations tested, the effect of depth was more profound than the effect of turbidity. Depth affected all parameters investigated except sponge size, whereas turbidity only affected specific pumping rates normalized to sponge surface area. Deep water sponges had clearly smaller oscula than shallow water sponges, but partially compensated for this lower pumping potential by showing a higher osculum density. Both increasing turbidity and increasing depth considerably decreased volumetric pumping rates of *C. reniformis*. These findings have important implications for selecting sites for mariculture of this species.

## 1. INTRODUCTION

Sponges (Porifera) are found at all latitudes, living in a wide array of ecosystems varying in temperature and depth (Hooper & Van Soest 2002; Van Soest et al. 2012). They are filter feeding organisms often dominating the benthos in abundance and biomass (Reiswig 1971; Reiswig 1975; Bell 2008; Ribes et al 2012). Efficient pumping of the surrounding seawater through the sponge body is of vital importance to sponges with regard to processes such as respiration and reproduction and feeding (Bergquist 1978; Simpson 1984). Sponges have a high efficiency and capacity for particle retention (Reiswig 1971, 1975, Ribes et al. 1999), preferably small particles (<10-µm), such as bacteria (Reiswig 1975b; Milanese et al. 2003; Fu et al. 2006; Zhang et al. 2010), phytoplankton (Reiswig 1971; Pile et al. 1996;) and even viruses (Hadas 2006). More recent, it has become evident that sponges can also utilize dissolved organic matter as major component of their daily diet (Yahel et al. 2003; De Goeij et al. 2008, Mueller et al. 2014). The efficient and versatile filtering makes sponges key drivers of the uptake, retention and transfer of energy and nutrients within benthic ecosystems (Gili and Coma 1998; Perea-Blázquez et al. 2012; De Goeij et al. 2013) and makes them interesting candidate species for bioremediation of organic pollution, such as the waste streams from aquacultures (Pronzato 1999; Milanese et al. 2003; Osinga et al. 2010). Recent studies reported successful trials of marine sponges as an environmental remediator of bacteria in integrated multi-trophic aquaculture (IMTA) designs. The sponge *Chondrilla nucula* retained up to 70 billion *E. coli* cells per square meter by filtering 14 L of water per hour (Milanese et al. 2003). A similar study showed a similar remediation of the bacteria *E. coli* and *Vibrio anguillarum* by the sponge *Hymeniacidon perleve* (Fu et al. 2006).

In order to fully comprehend the role of sponges in transfer of nutrients and energy in natural and artificial ecosystems such as IMTAs, it is important to understand the mechanism of the sponge pump and how it is affected by physiological and environmental factors. Sponges actively pump water through their filter system, termed ‘aquiferous system’, a maze of canals and filter chambers, lined by flagellated filter cells, the choanocytes (Simpson 1984). Water enters the sponge through incurrent pores termed ostia and leaves the sponge through larger exhalant openings, the oscula (Bergquist 1978). The size and morphology of the oscula varies among species (Vogel 1974; Reiswig 1975). The outflow velocity of the exhalant flow has to be sufficient to prevent re-inhaling the processed water (Riisgard et al. 1993).

Sponge pumping and filtration capacities have been studied and quantified extensively, both *in situ* and in laboratory studies (Reiswig 1971, 1974; Vogel 1974, Riisgard et al. 1993; Mendola et al. 2008). Sponges process huge amounts of water daily, up to 50,000 L seawater per L sponge volume per day (Weisz et al. 2008), which is comparable to well-established suspension feeders, such as mussels (Jorgensen 1949, 1954; Riisgard and Larsen 2000). Limited knowledge is available on possible factors that control *in situ* pumping activities of sponges. Sponges hosting large quantities of associated microbes often have lower pumping rates than sponges with low numbers of associated microbes (Weisz et al. 2008). Additionally, ambient flow may influence sponge pumping activity by reducing the amount of energy that is needed for active pumping (Vogel 1974). Increased turbidity, the concentration of suspended matter in water, can have a negative effect on sponge pumping activities (Reiswig 1971a; Mendola 2008). High (in)organic loading can lead to suffocation of sponges and consequently, to an arrest in pumping. Gerrodette & Flechsig (1979) reported that elevated amounts of suspended sediment (various sediment concentrations-natural marine clay of quartz, illite and chlorite) in the surrounding water reduced the pumping activity of sponges. Similarly, the filtration rate of the sponge species *Pseudosuberites aff. andrewsi* was affected considerably by the concentration of unicellular algae (*Dunaliella*) in the surrounding water (Osinga et al. 2001).

In this study we describe the effects of turbidity and depth on *in situ* sponge pumping activity of the sponge *Chondrosia reniformis* (Nardo, 1847). *C. reniformis* is a frequently occurring sponge species on rocky substrates at a depth of 1 to 50 m along the Eastern Mediterranean coastline, where it often dominates the benthic cover (Wilkinson and Vacelet 1979; Parma et al. 2007; Gokalp 2012). This ubiquitous species has also economical relevance, as producer of collagen for applications in human therapy (Swatschek et al. 2002; Nickel and Brummer 2003). Hence, knowledge on the pumping activity of this species is relevant to benthic ecology and to aquaculture.

## 2. MATERIALS AND METHODS

### 2.1 Model species, study area and site selection

Field studies were conducted along the southwestern coastline of Turkey, around the Bodrum Peninsula (Southwest Turkey, Fig. 1) in May-December 2012. The sub-littoral rocky coastline of Bodrum Peninsula provides ideal habitat for *C. reniformis*, which prefers shady habitats such as the vertical sides or bottom sides of rocks, crevices, overhangs and caves, although it is occasionally also found on sandy bottoms and in *Posidonia* meadows.

**Fig. 1.**
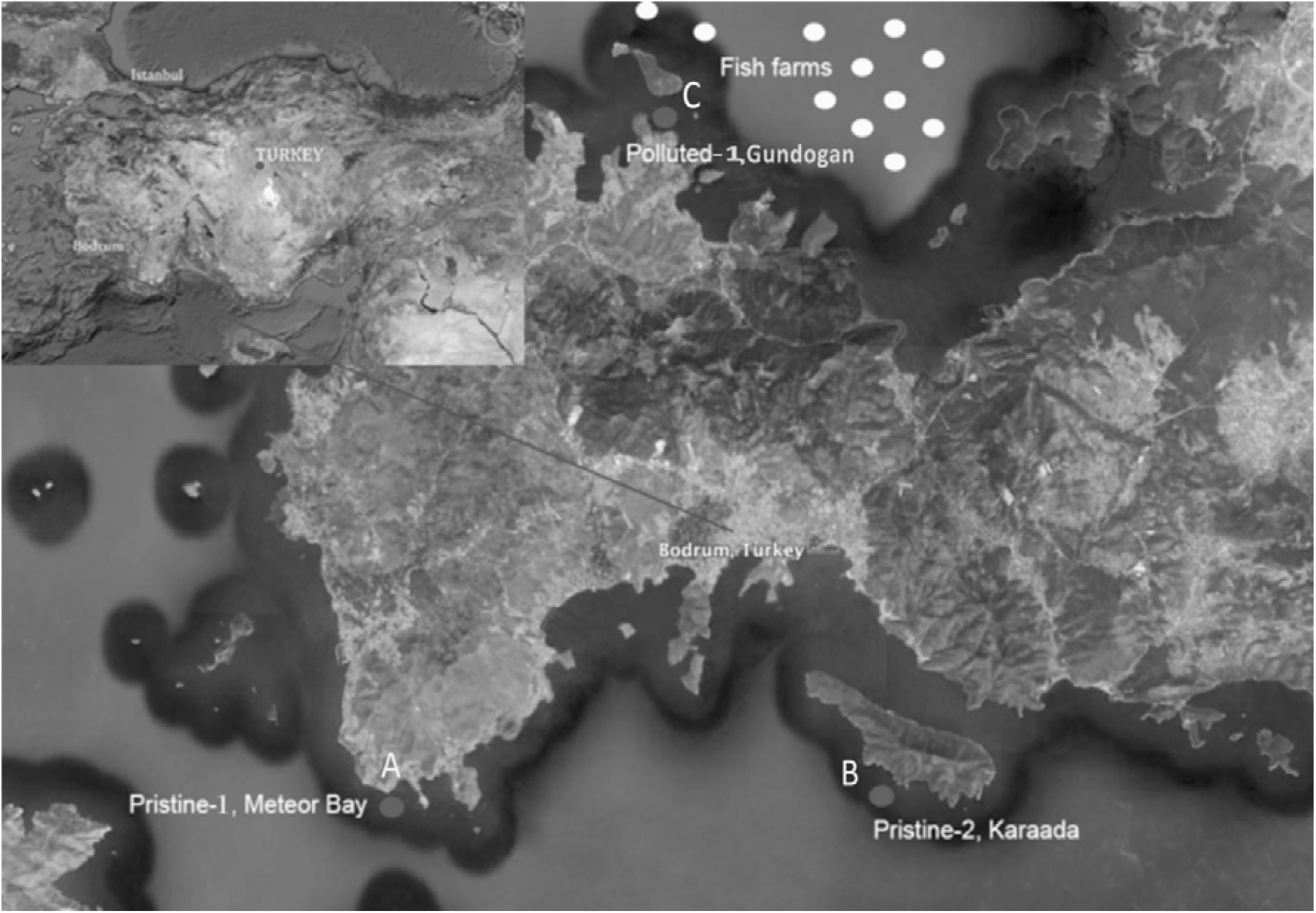
Map of Bodrum Peninsula: Polluted site located at the northern side of the Bodrum Peninsula along with intensive fish farming activities in the close proximity. Two pristine sites located at the southern side of the Bodrum Peninsula with clear waters. Global positioning system coordinates of the three locations; **A**-36°57’50. 33”N, 27°16’35.11”E **B**-36°58’6.64”N, 27°27’2.59”E, **C**-37°9’8.14”N, 27°22’9.60”E.

The Bodrum peninsula waters span different habitats, ranging from pristine areas to heavily polluted sites that are strongly influenced by mariculture waste products. Therefore, this region represents a suitable location for measurements of *in situ* sponge pumping under different levels of turbidity. The southern side of the peninsula represents a good example of a pristine, oligotrophic environment, which receives daily replenishment from deeper waters through a strong, continuous oceanic current termed “Kos current”. Conversely, the northern side shows characteristics of a secluded bay. This area is eutrophic, mainly as a result of intensive open water fish farming activities (Demirak et al. 2006; Basaran et al. 2010). By doing a SCUBA survey, three specific study sites (marked A, B and C in Fig. 1) were selected that were considered suitable for comparative studies on the effects of depth and turbidity on sponge pumping behavior. It should be noted that no specimens of *C. reniformis* were found in the deeper, muddy parts of the polluted bay.

*Site A (Meteor Bay)* is a shallow pristine site at the southern side of the peninsula (0-3 m). The site is receiving daily fresh waters from the current in between Kos Island and Bodrum Peninsula. It is a cove, almost completely protected (except when southerly winds occur) from wave action and high currents, exhibiting nearly still waters during the summer period. However, during winter, spring and autumn periods, wave action and intermediate storms are regular events. The bottom is mostly rocky with irregular *Posidonia oceanica* meadows. At this location, patches of *C. reniformis* are mostly found attached on sides or lower portions of the rocks.

*Site B (Karaada)* is a deep pristine site at the southern side of the peninsula (20-25 m), south of Karaada Island. It is a rocky shore at the tip of a steep inclined hill receiving wave action and water flow irregularly (Fig. 2b). At this location, large boulders and rocks are covered on all sides with healthy *C. reniformis* colonies.

**Fig. 2.**
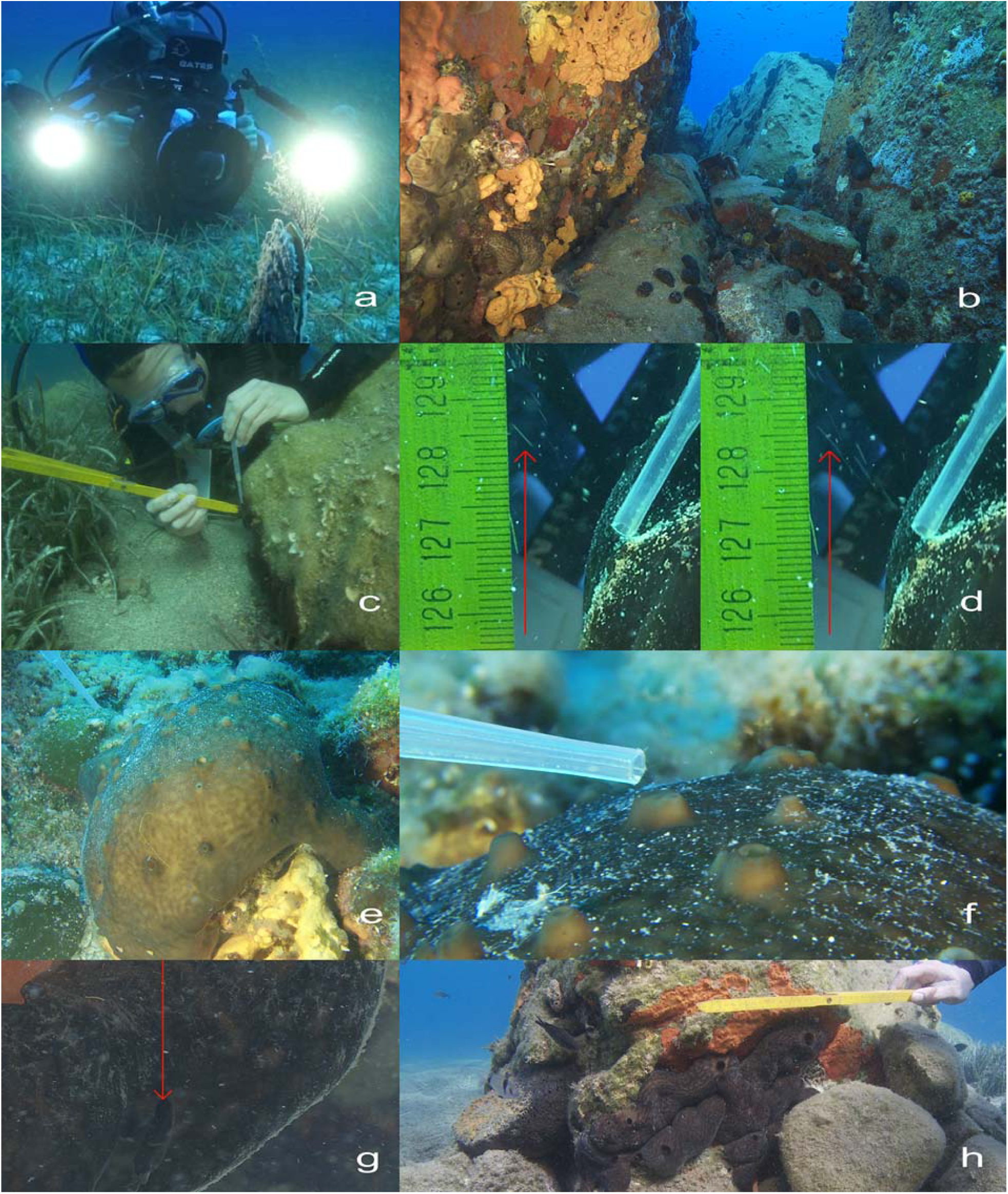
Oscular flow velocity measurements **a** Underwater setup. **b** Pristine deep site, (visibility close to 30 meters). The healthy colonies of *C. reniformis* are located on the side of the walls and at the bottom along with other Mediterranean sponge species. **c** Polluted site, the diver is ready to releases neutrally buoyant particles over the oscula in the meantime the other diver positions the HD video device perpendicular to the target oscula. **d** Pumping sponge oscula & two consecutive frames of a video clip. The trajectories of the particles are shown with arrows. **e.** Pristine deep site, deep specimen of *C. reniformis* with many small sized osculas. **f** A closer shot of the same specimen (picture e), the oscular sizes ranges from 0.05 to 0.2 cm. **g** Pristine shallow site, a *C. reniformis* specimen with large single osculum. **h** On-site measurements of oscular size, total oscular number per sponge and sponge patch size.

*Site C (Gundogan)* is a shallow polluted site at the northern part of the peninsula (2-10 m). The site was selected because of its proximity to fish farming activities and the observed presence of *C. reniformis* specimens with healthy outer appearance. It is a shallow area, relatively less polluted than the inner portions of the adjacent Guvercinlik Bay and composed of small and medium sized rocks surrounded with *P. oceanica* meadows. The seafloor at this site is subject to wave action that varies in strength depending on the weather and wind direction. At this site, *C. reniformis* specimens are found attached on the sides or on the lower portions of rocks and stones.

### 2.2 Measurement methods

Exhalant flow velocities were measured by tracking the particle movement in the exhalant stream of selected osculum (Fig. 2d). The monitoring was performed by two SCUBA divers, where one diver positioned an HD video device perpendicular to the targeted osculum (Fig. 2c), while the other diver released nearly neutrally buoyant particles into the exhalant stream directly above the sponge osculum using a (0.2 mm) pipette (Fig. 2f). Detrital materials sampled from the surface of other sponges were used for this purpose. The impulse of the exhalant flow accelerated the particles and gave them a particular speed. The trajectories of the particles travelling with the efflux were recorded with a video camera (Sony PMW Ex-1 HD video camera, Gates Ex-1 housing, Gates, macro port, Green Force 250 HID video light and TIII battery setup) attached to a tripod head (Fig. 2a). The video recordings were made at 1280×720 HD 16:9, recording speed was set at 60 frames per second at all measurements. Only particles with straight trajectories that depart the center of the osculum were taken into account. The slower-exiting particles that were released nearer to the walls of the oscula were discarded. The distance between the lens and subject was minimized to increase spatial resolution and particle recognition. A ruler was placed behind the recorded sponge osculum as a reference at all times. Additionally, osculum sizes, total oscular number per sponge and sponge surface area were measured with a ruler and documented on site for all sponge specimens selected for oscular outflow velocity analysis (Fig. 2h).

The time coded videos were transferred to Final cut Pro editing software via XDCAM transfer program. The outflow velocities were calculated by analyzing the trajectory of the buoyant particles using frame-by-frame video analysis after the principles of the technique of Savarese et al. (1997). There is 0.016677 seconds in between each frame at 60 fps video recording speed. Hence, the difference in position of the particle relative to the ruler between two subsequent frames equals the distance travelled by the particle in 0.0167 seconds. Dividing the change in position relative to the ruler between two subsequent frames by 0.0167 s gave us the oscular outflow velocity in cm s^-1^. The distance travelled by one particle within a particular time frame was measured using the picture editing software Image Tool. The movement of several particles leaving the oscular opening was averaged together to obtain a velocity for each observation in between the two subsequent frames picturing particle at its maximum velocities. Using the osculum diameters, outflow velocities (in cm s^-1^) were multiplied with the oscular area to obtain a pumping rate (in cm^3^ s^-1^) by using the following equation below:

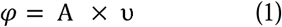

in which *φ* is the pumping rate (cm^3^ s^-1^), ν is the oscular outflow velocity (cm s^-1^), and A is the oscular surface area (cm^2^).

In order to minimize effects of daily rhythms on pumping activity, every recording was taken between 9 am and 1 pm, hereby taking into account Reiswig’s (1971) earlier observations on variability of pumping activity of sponges, where maximum pumping activity occurred when water circulation over the habitat was maximal.

### 2.3 Temperature and Turbidity Measurements

Water temperature (Uwatec Aladin Air X Nitrox dive computer) and visibility (Secchi disk, cf. Hannah et al., 2013) were measured 17 times during periodical visits at the culture platforms throughout the experimental period. To assess the degree of organic pollution, three replicate water samples were taken within 10 m from the culture platforms by scuba diving from each location for analysis of Total Organic Carbon using a wet oxidation method (Menzel & Vaccaro, 1964). The seawater samples were collected in August 2015, and were stored in pre-combusted 50 ml glass bottles at -20 C° until analysis. Prior to analysis, sulphuric acid was added to the samples (end concentration 0.002 M) to remove dissolved inorganic carbon species. The acidified samples were diluted 5 times with distilled water, supplemented with sodium tetraborate and potassium persulphate and processed using segmented flow analysis (SFA) on a Continuous Flow Analyser (Skalar, Breda, The Netherlands). In SFA, TOC is first oxidized using UV light and then measured as CO2 using infrared detection.

### 2.4 Characterization of pumping behavior

Pumping activity of *C. reniformis* was studied in two consecutive steps. First, it was determined how oscular outflow and osculum diameter correlated under four different combinations of turbidity and depth (Table 2). Oscular outflow and corresponding osculum diameter were measured as described above for at least 15 oscula per turbidity/depth combination (Table 2). These measurements were done during a first survey in May 2012. Second, two comparative studies were executed between June 2012 and August 2012 in order to separately investigate the effect of turbidity and the effect of depth on the pumping behavior of *C. reniformis*. For these studies, measurements of oscular diameter, oscular numbers per sponge specimen and surface area per sponge specimen were done in three different types of environments: Pristine/Shallow, Pristine/Deep and Polluted/Shallow, on a variable number of sponges per environment (see Table 2 for details). Specimens were highly variable in size, and it was sometimes difficult to distinguish between individuals. We included only individuals that were clearly separated from others. The environment type Polluted/Deep was not included, since no specimen of *C. reniformis* were be found in the deeper, muddy parts of the polluted area. Hence, we compared Pristine/Shallow to Pristine/Deep to investigate the effect of depth and we compared Pristine/Shallow to Polluted/Shallow to investigate the effect of turbidity.

**Table 1.** Overview of measurements done to investigate the relation between oscular size and oscular outflow velocity under different environmental conditions.

| Site | Number of oscula<br>investigated | Depth range<br>(m) |
| --- | --- | --- |
| A – shallow/pristine | 30 | 2-3 |
| B – deep/pristine | 38 | 20-25 |
| C1–shallow/polluted | 43 | 2-3 |
| C1– deep/polluted | 15 | 7-15 |

**Table 2.** Overview of the number of measurements applied for the comparative studies on depth and turbidity.

| Depth comparison |  |  |  |
| --- | --- | --- | --- |
| Site | Depth (m) | Number of sponge individuals investigated | Number of sponge oscula investigated |
| A – shallow/pristine | 0-3 | 15 | 87 |
| B – deep/pristine | > 20 | 10 | 298 |
| Turbidity comparison |  |  |  |
| Site | Depth (m) | Number of sponge individuals investigated | Number of sponge oscula investigated |
| A – pristine | 0-10 | 25 | 222 |
| C – polluted | 0-10 | 21 | 174 |

To further characterize pumping activity, a series of additional characteristics (Table 3) was deduced from the data obtained during the second survey for osculum diameter, osculum numbers per sponge and sponge surface areas, hereby using the data obtained during the first measurement series to calculate oscular outflow velocity from osculum diameter. In case a significant correlation between diameter and outflow velocity had been found for a particular site, a linear regression equation was used to calculate outflow velocity from diameter, in case the correlation was not significant, the average outflow rate was used for all oscula to calculate outflow velocity from osculum diameter. Using this calculated approximation of oscular outflow, the total pumping rate and specific pumping rate per individual sponge was calculated with equations (1) and (2):

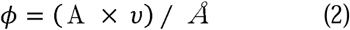

in which *ϕ* is the specific pumping rate (cm^3^ cm^-2^ s^-1^), *Å* is the surface area of the individual sponge patch (cm^2^).

**Table 3.** Overview of calculations deduced from the measurements of the osculum diameter, sponge surface area and total number of oscula per sponge individual.

| Parameter determined | Unit | Method / calculation |
| --- | --- | --- |
| Osculum size | cm<sup>2</sup> | $\pi(\text{Diameter}/2)^2$ (assuming circular shape) |
| Average osculum size | cm<sup>2</sup> | $\Sigma(\text{osculum size})/\text{number of oscula measured}$ |
| Osculum density – the number of oscula per cm<sup>2</sup> of sponge surface area | cm<sup>-2</sup> | Counts divided by sponge surface area |
| The ratio between osculum size and sponge surface area | cm<sup>2</sup> cm<sup>-2</sup> | $\Sigma(\text{osculum size}) / \text{sponge surface area}$ |
| Individual pumping rate per osculum | cm<sup>3</sup>s<sup>-1</sup> | $\Sigma(\text{osculum size} \times \text{oscular flow})$ |
| Pumping rate per unit sponge surface area (specific pumping rate) | cm<sup>3</sup> cm<sup>-2</sup> s<sup>-1</sup> | $\Sigma(\text{osculum size} \times \text{oscular flow}) / \text{sponge surface area}$ |

### 2.5 Statistical analysis

The collected data for osculum diameters and corresponding outflow velocities (the first measurement series in May 2012) were split in “pristine” and “polluted” groups according to the water turbidity level of the sponge habitat. The data of both groups were separated into two distinct sub-groups (Shallow 2-3 m & Deep > 10 meters) according to the measurement depth of the sponge oscula and checked for normal distribution (Kolmogorov-Smirnov test). Depending on the result of this test, either Pearson correlation analysis or the non-parametric Spearman correlation analysis was applied to relate oscular size to oscular outflow velocity for each location. Data for oscular diameter, oscular numbers per sponge and sponge surface area obtained during the second series of measurements (June-August 2012) were compared between the two depth classes and the two turbidity levels, as indicated in Table 2. The data of each category were tested for normal distribution (Kolmogorov-Smirnov test). Depending on the results, Students t-test was applied in case of normality and a Mann-Whitney test was applied in case of non-normal data. The comparison was done for all parameters listed in Table 3.

## 3. RESULTS

During the period of this study, we measured Secchi disk depths varying in between 4 - 9 m at the shallow portions of Guvercinlik Bay and 8-14 m at the Gundogan site, northern side of Bodrum Peninsula. Our Secchi disk recordings at the pristine shallow and deep sites varied in between 18-24 m and 20-30 m. respectively. Comparable Secchi depths have been reported by local institutions (e.g. Bingel et. al 2005; Eryalcin et al. 2007). Secchi disk depth measurements varying in between 25-32 m was reported for the deeper portions of the southern side of Bodrum Peninsula, whereas at the northern side, the recordings decreased from 29 m to 10 m approaching from offshore to the inner parts of the bay. In 2006-2007 around Salih Island where extensive aquaculture activities were taking place, Eryalcin et. al (2007) reported Secchi disk depths as low as 3 to 6 m. Our measurements indicated that, inside Guvercinlik bay TOC levels as high as 2.5 times was recorded compared to the pristine site (Mean TOC _polluted, pristine_ = 3.34 ± 0.86, 1.37 ± 0.08 mg C/L, respectively). Moreover, Secchi depth measurements (cf. Hannah et al., 2013) during this study revealed 4 times higher visibility in the pristine site compared to the polluted site (Mean Sechhi Disk _polluted, pristine_ = 6.5, 25 meter, respectively), which is an earlier fish farm site. The water temperatures for both sites didn’t differ, measuring in between 13 - 27° C.

### 3.1 Correlation between oscular size and outflow velocity

Osculum diameters and corresponding rates of oscular outflow velocity measured at four different environmental conditions are shown in Fig. 3. Overall, osculum diameter ranged between 0.05 and 2.0 cm (mean 0.53 ± 0.43 cm). Outflow velocities ranged between 2.5 to 12.5 cm s^-1^ with a mean velocity of 6.81 ± 2.86 cm s^-1^. None of the four data series obtained during the first survey for osculum size and oscular outflow were normally distributed. Hence, Spearman rank correlation was applied to test whether relations between the two parameters were significant. The relationship between oscular diameter and the oscular outflow velocity at the pristine deep location was found to be insignificant Pristine/Deep (Spearman rank correlation rs = 0.324, N=36, *P* (1-tailed) > 0.001, Fig. 3b). Oppositely, significant positive relationships between oscular diameter and the oscular outflow velocity were found for Pristine/Shallow (Spearman rank correlation = 0.678, N= 30, *P* (1-tailed) < 0.001, Fig. 3a), Polluted/Shallow (Spearman rank correlation *rs* = 0.623, N=43, *P* (1-tailed) < 0.001, Fig. 3c) and Polluted/Deep (Spearman rank correlation *rs* = 0.882, N=15, *P* (1-tailed) < 0.001, Fig. 3d). The linear relations of the trendlines for these three environment types are given by the equations 3-5.

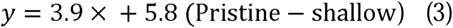

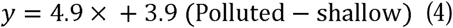

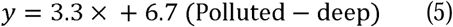

**Fig. 3.**
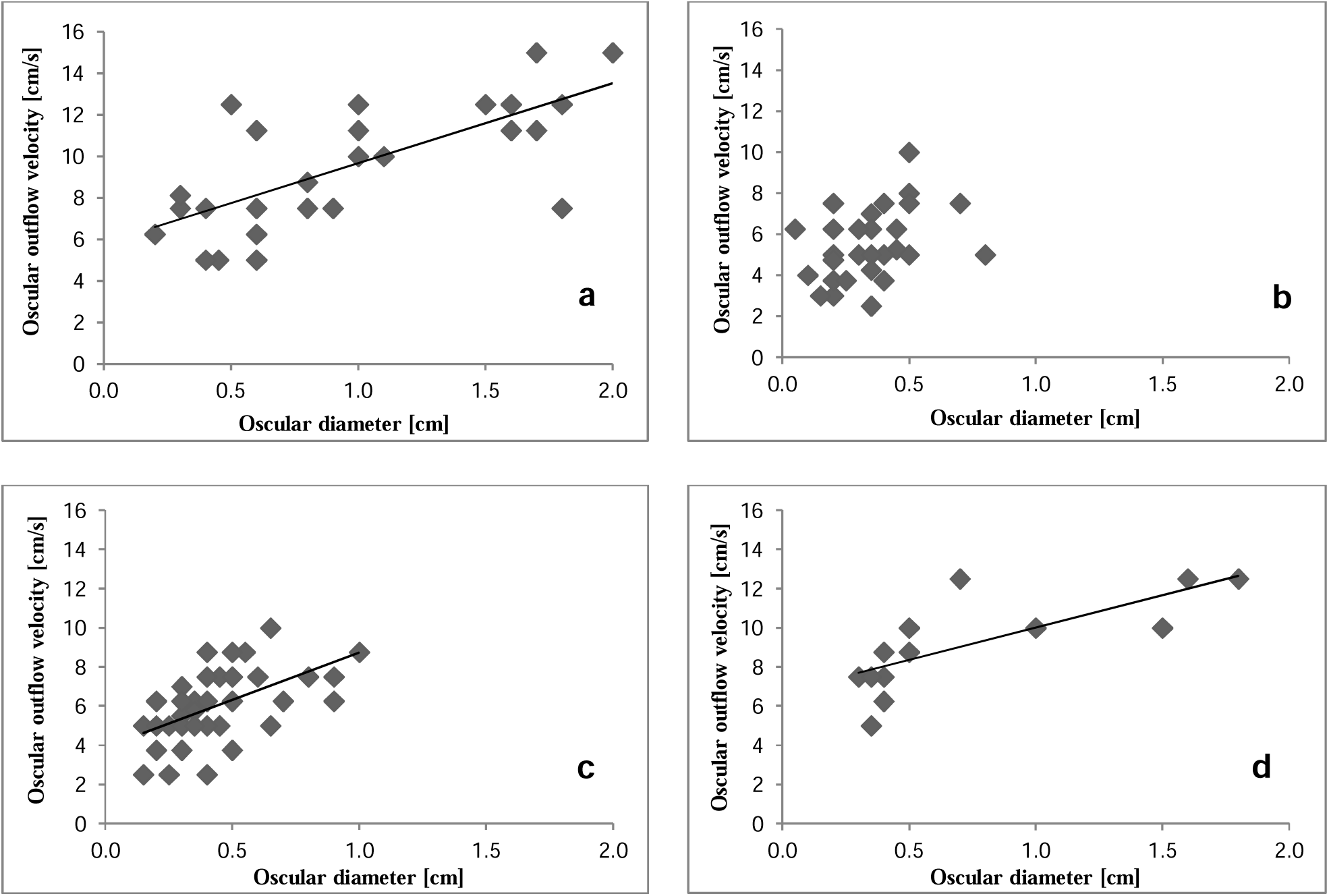
Correlations between pairs of variables analyzed **a** Correlation between average osculum size and oscular outflow velocity at the pristine shallow site **b** pristine deep site **c** polluted shallow site **d** polluted deep site

The outflow velocities for the oscula documented during the second survey were calculated by using the equations 3-5 above (i.e. not including Pristine/Deep, for which the average outflow velocity was used) and subsequently, total pumping rate and pumping rate per sponge surface area were calculated using Equations 1 and 2.

### 3.2 Effect of turbidity

The results of the comparisons between *C. reniformis* specimens monitored at two different turbidity conditions (0 - 10 m depth range) can be seen in Table 4. Sponges inhabiting the pristine locations had an osculum density ranging from 0.01 to 0.41 oscula per cm^2^ of sponge (median = 0.05), whereas at the polluted location, osculum density was found to be within a range of 0.02 - 0.64 oscula cm^-2^ (median = 0.07). The difference in osculum density among the sponges living in two different water turbidity conditions was statistically insignificant (Mann–Whitney *z* = -0.02, N1=25, N2=21, *P* > 0.05). Mean osculum size ranged from 0.01 – 2.54 cm^2^ (median = 0.45 cm^2^) under pristine conditions and from 0.02 – 0.79 cm^2^ (median = 0.13 cm^2^) under polluted conditions. Despite the three-fold difference in medians, the difference between the two groups was found to be insignificant (Mann–Whitney *z* = -0.88, N1=38,N2=66, P > 0.05). The difference in the total surface area of the sponges was also insignificant (Mann–Whitney *z* = -0.03, N1=25, 21, *P* > 0.05) as well as the ratio between the total osculum surface area and the corresponding sponge surface area (Mann– Whitney *z* = -0.441, N1=25, N2=21, *P* > 0.05). In contrast, the average individual oscular pumping rates calculated for pristine and polluted conditions (median _pristine, polluted_ = 2.45, 0.63 cm^3^s^-1^ respectively) revealed a significant difference (Mann–Whitney, *z* = -2.71, N1=25, N2=21, *P* < 0.05). The specific pumping rate per cm^2^ of sponge surface was found to be more than 3 fold higher for pristine conditions than for polluted conditions (0.28 and 0.08 cm^3^ per cm^2^ sponge area per second, respectively) and this difference was also significant (Mann–Whitney *z* = -2.09, N1=25, N2=21, *P* < 0.05).

**Table 4.** The medians, averages, ranges and the level of significance (Mann–Whitney test) of the different sponge characteristics ar e summarized for *C. reniformis* specimens that were monitored at two different turbidity conditions.

| Sponge Characteristics | Range |  | Median |  | Average |  | Significance<br>(*P*-Value) |
| --- | --- | --- | --- | --- | --- | --- | --- |
|  | Pristine | Polluted | Pristine | Polluted | Pristine | Polluted |  |
| Sponge surface area (cm<sup>2</sup>) | 15-520 | 10-590 | 72 | 80 | 118 ± 136 | 129 ± 142 | 0.5 |
| Average osculum size (cm<sup>2</sup>) | 0.01-2.54 | 0.02-0.79 | 0.45 | 0.13 | 0.49 ± 0.63 | 0.20 ± 0.22 | 0.4 |
| Osculum density (cm<sup>-2</sup>) | 0.01-0.41 | 0.02-0.64 | 0.05 | 0.07 | 0.15 ± 0.12 | 0.11 ± 0.13 | 0.5 |
| Ratio oscular area per sponge area (cm<sup>2</sup>cm<sup>-2</sup>) | 0.002-0.141 | 0.001-0.039 | 0.017 | 0.011 | 0.02 ± 0.03 | 0.01 ± 0.08 | 0.67 |
| Individual pumping rate per osculum (cm<sup>3</sup>s<sup>-1</sup>) | 0.02-47.1 | 0.04-18 | 2.45 | 0.63 | 7.9 ± 11.1 | 1.9 ± 3.4 | 0.006 |
| Specific pumping rate per cm<sup>2</sup> sponge (cm<sup>3</sup>cm<sup>-2</sup>s<sup>-1</sup>) | 0.01-1.80 | 0.01-0.34 | 0.19 | 0.06 | 0.28 ± 0.40 | 0.08 ± 0.1 | 0.036 |

### 3.3 Effect of depth on pumping

The results of the comparison between *C. reniformis* specimens monitored at two different depth levels can be seen in Table 5. The osculum density for sponges inhabiting the shallow zone (0.01 cm to 0.27 oscula per cm^2^ sponge area; median = 0.04 cm^-2^) was significantly lower (Mann–Whitney *z* = -3.41, N1=17, N2=10, *P* < 0.05) than for sponge in the deep zone (0.1 cm to 0.83 oscula per cm^2^ sponge area; median = 0.24 cm^-2^). Average osculum size in shallow water (0.71 ± 0.67 cm^2^), however, was significantly higher (Student’s T-Test *z* = -4.1, N1=31, N2=40, *P* <0.05) than average osculum size in deep water (0.05 ± 0.02 cm^2^). The ratio between osculum surface area and sponge surface area at the two different depth zones (median _shallow, deep_ = 0.02, 0.01 respectively) also differed significantly (Mann–Whitney *t* = -3.6, N1=17, N2=10, *P* <0.05), the shallow sponges had three times more osculum area per cm^2^ sponge surface. The calculated pumping rates per osculum recorded at the shallow and deep zones ranged from 0.20 to 47.12 cm^3^ s^-1^ and from 0.01 to 2.89 cm^3^ s^-1^, respectively, revealing another significant difference (Mann–Whitney, *z* = -5.95, N1=17, N2=10, *P* < 0.05) between the two depth zones. The specific pumping rate per cm^2^ of sponge surface (Median _shallow, deep_ = 0.17, 0.06 respectively) was found to be significantly different between the two depth zones (Mann–Whitney, *z* = -3.41, N1=17, N2=10, *P* < 0.05). The sponges were not significantly different in size at the two different depth zones (Student’s T-Test, *t* = -0.363, N1=17, N2=10, *P* > 0.05), (Average _shallow, deep_ = 130, 146 cm^2^, respectively).

### 3.4 Average volumetric pumping activity of *C. reniformis*

In order to convert the data on pumping per cm^2^ of sponge area to volumetric pumping rates, an average tissue thickness of 2.3 cm was assumed for *C. reniformis* (based upon a series of random measurements of sponge thickness). All surface related data were divided by 2.3 to obtain volumetric pumping activity. The overall range of volumetric pumping by *C. reniformis* in our study (i.e. regardless of location) was thus estimated at 0.003-0.64 cm^3^ (cm^3^ sponge)^-1^ s^-1^.

## 4. DISCUSSION

Pumping activity of *C. reniformis* – the role of osculum size

Osculum diameters and oscular outflow velocities measured in this study are within the range reported for other sponge species (Table 6). The range of outflow velocities of 2.5 – 12.5 cm s^-1^ determined for *C. reniformis* is quite close to Mendola’s (2008) findings with another Mediterranean species, *Dysidea avara* (3.5 – 13 cm s^-1^), which has similarly sized oscula. Reiswig (1971, 1974, 1975a), who substantiated the most extensive sponge pumping activity study to date, reported comparable outflow velocities ranging in between 2.5 - 18.6 cm s^-1^ with Caribbean sponges. We conclude that our method for determining oscular outflow velocity is accurate and provides a good alternative for previously used methods.

Overall, the *C. reniformis* specimen tested in this study showed a significant positive relation between the oscular diameter and the outflow velocity of the water exiting the sponge osculum. This observation substantiates the observations of Reiswig (1971) on three tropical *Demosponge* species *Mycale sp*., *Tethya crypta* and *Verongia gigantea* that bigger oscula exhibit higher pumping rates per unit of osculum surface. Bigger oscula are apparently more efficient than smaller oscula: they do not only process more water absolutely, but also relatively.

For *C. reniformis*, the strength of the relation between size and outflow differed among localities. Hence, one cannot apply a general ratio between the two parameters to deduce velocity from size; this is only possible on a site-by-site basis. It also seems that the relationship is not linear, but levels off when oscula get bigger. The exact nature of this relationship needs to be further substantiated by studying larger number of oscula.

The range of calculated volumetric water process rates for *C. reniformis* is in good agreement with literature data for other species (e.g. Weisz et al. 2008) and spans nearly the entire range of the previously reported rates. The highest rate recorded in this study (0.64 cm^3^ cm^-3^ s^-1^) is among the highest rates ever recorded for sponges. This rate was calculated for a specimen with a single, giant osculum that had a diameter of 2 cm, which confirms that sponges exhibiting large oscula are more efficient pumpers than their conspecifics exhibiting smaller oscula.

**Table 5.** The medians and averages (for the groups analyzed with the non – parametric Mann–Whitney test & Student’s T-TEST) and ranges of the different sponge characteristics are summarized for *C. reniformis* specimens that were monitored at the pristine locations at two different depth levels.

| Sponge Characteristics | Range |  | Median |  | Average |  | Significance<br>(*P*-Value) |
| --- | --- | --- | --- | --- | --- | --- | --- |
|  | Shallow<br>(0-3 m) | Deep<br>(20-30 m) | Shallow<br>(0-3 m) | Deep<br>(20-30 m) | Shallow<br>(0-3 m) | Deep<br>(20-30 m) |  |
| Sponge surface area (cm<sup>2</sup>) | 15-520 | 23-290 | 85 | 153 | 130±147 | 146±83 | 0.72 (T-Test) |
| Average osculum size (cm<sup>2</sup>) | 0.07-2.54 | 0.03-0.1 | 0.5 | 0.04 | 0.71±0.67 | 0.05±0.02 | 0.001 (T-Test) |
| Osculum density (cm<sup>-2</sup>) | 0.01-0.27 | 0.1-0.83 | 0.04 | 0.24 | 0.07 ± 0.08 | 0.27 ± 0.2 | 0.000 (MW) |
| Ratio oscular area per sponge<br>area (cm<sup>2</sup>cm<sup>-2</sup>) | 0.011-0.141 | 0.004-0.027 | 0.02 | 0.01 | 0.03 ± 0.03 | 0.01 ± 0.01 | 0.000 (MW) |
| Individual pumping rate per<br>osculum (cm<sup>3</sup>s<sup>-1</sup>) | 0.2-47.1 | 0.01-2.9 | 4.4 | 0.4 | 9.9 ± 11.8 | 0.6 ± 0.7 | 0.000 (MW) |
| Specific pumping rate per cm<sup>2</sup><br>sponge (cm<sup>3</sup>cm<sup>-2</sup>s<sup>-1</sup>) | 0.1-1.9 | 0.02-0.17 | 0.17 | 0.06 | 0.29 ± 0.43 | 0.06 ±0.04 | 0.000 (MW) |

**Table 6.** Comparison of the previously published sponge pumping activity data in the literature

| Sponge Species | Oscular Size<br>(cm) | Oscular outflow speed<br>(cm/s) | Pumping rate<br>$\text{cm}^3 (\text{cm}^3 \text{ sponge})^{-1} \text{s}^{-1}$ | Reference |
| --- | --- | --- | --- | --- |
| *Spinosella sororia* |  | 4 | 0.010 | (Parker 1910, 1914) |
| *Leucandra aspera* |  | 7.0 - 8.5 | 0.026 | (Bidder 1923) |
| 8 species (grouped) |  | 7.5 - 22 | 0.20– 0.40 | (Vogel 1974) |
| *Aplysina fistularis* |  |  | 0.049–0.276 | (Reiswig 1974) |
| *Verongia gigantea* |  | 2.5 – 16.8 | 0.040–0.15 | (Reiswig 1971, Reiswig 1974) |
| *Tethya crypta* | > 10 | 5.5 – 18.2 | 0.11 – 0.22 | (Reiswig 1971, Reiswig 1974) |
| *Mycale* sp. | 2 - 10 | 6.6 – 9.1 | 0.210–0.340 | (Reiswig 1971, Reiswig 1974) |
| *Haliclona permollis* | 0.14 | 8.5 – 9.2 | 0.049 – 0.131 | (Reiswig 1975a) |
| *Aplysina lacunosa* |  |  | 0.005-0.034 | (Gerrodette & Flehsig 1979) |
| *Verongia fistularis* | 3.3 | 8.6 - 11.6 | 0.049-0.276 | (Reiswig 1981) |
| *Haliclona urceolus* |  | 5.4 | 0.042 | (Riisgard et al. 1993) |
| *Halichondria panicea* | 0.015 | 5.6 | 0.045 | (Riisgard et al. 1993) |
| *Mycale lingua* | 1.3 – 2.3 | 14 |  | (Pile et al. 1996) |
| *Thenea muricata* |  |  | 0.05 | (Witte et al. 1997) |
| *Dysidea avara* |  |  | 0.017 | (Ribes et al. 1999) |
| *Baikalospongia bacillifera* |  | 0.2 – 3.3 | 0.040 – 0.60 | (Savarese et al. 1997) |
| *Spechiospongia vesperium* |  |  | 0.069 | (Lynch & Philips 2000) |
| *Halicondria melanadocia* |  |  | 0.080 | (Lynch & Philips 2000) |
| *Theonella swinhoei* | 0.6 -1.5 | 7.6 | 0.043 | (Yahel et al. 2003) |
| *Dysidea avara* |  | 3.5 - 13 | 0.008 – 0.30 | (Mendola 2008) |
| 8 species (grouped) |  |  | 0.073 – 0.578 | (Weisz et al. 2008) |
| *Aphrocallistes vastus* | 7 | 0.5 – 5.2 | 0.05 | (Leys et al. 2011) |
| *Chondrosia reniformis* | 0.05 - 2 | 2.5 – 12.5 | 0.01 – 0.64 | This study |

### 4.1 Turbidity Effect

Total sponge surface area, osculum size, osculum density and total osculum area per sponge area did not differ significantly between the pristine location and the polluted location. Despite these similar characteristics of the pumping apparatus, the conditions at the polluted (i.e. more turbid) location considerably suppressed pumping efficiency. Specific pumping rates per cm^2^ of sponge surface were over three times higher under pristine conditions. This finding is in agreement with earlier studies (Gerrodette & Flechsig 1979; Osinga et al. 2001; Mendola 2008), in which higher particle densities decreased sponge pumping rates. Osinga et al. (2001) described a tipping point for pumping of *Pseudosuberites aff. andrewsi*, which rapidly declined when particle density reached a particular threshold value. If pumping activity of *C. reniformis* would be related to turbidity in a similar way, the three-fold decrease in specific pumping rate may indicate that the conditions at the polluted site in this study are close to the upper limit of turbidity that can be sustained by this species. It is also possible that sponges do not pump as much in “polluted” sites because they don’t have to, to obtain the same amount of food. That is the polluted site may have higher food abundance.

### 4.2 Depth Effect

Water depth had a profound effect on the pumping system of *C. reniformis* and its activity. Sponges in the deep zone had more, but smaller oscula than sponges in the shallow zone. Hence, the sponges compensate for a decrease in average osculum size by building more oscula. Despite this compensation, shallow water sponges on average had twice as much osculum surface per unit of sponge surface. Moreover, sponges having larger average osculum size (i.e. the shallow water sponges in this study) process more water, due to the higher pumping rate per unit of osculum area exhibited by larger oscula. As a result, specific pumping rates per cm^2^ of sponge surface were nearly five-fold higher for *C. reniformis* specimen living in shallow water than for conspecifics living in deep water.

It remains open for discussion why sponges vary the size of their oscula and why deep water sponges have smaller oscula than shallow water sponges. Having large, fast pumping oscula will be more costly for the sponges in terms of metabolic energy requirements, since the pressure needed inside the sponge body to generate an exhalant current will increase when oscula become bigger. However, investing in a higher pumping potential may have several advantages for the sponges. First, having large oscula could be a physiological response to low food availability, i.e. when high amounts of water need to be processed to obtain sufficient nutrition. Nevertheless, the fact that we also found some very large oscula on sponges living in the deeper parts (7-10 m) of the polluted area does not support this explanation. Second, large oscula may represent a physiological adaptation to sedimentation. In shallow areas such as Meteor Bay, sand particles may resuspend as a result of wave action during storms. These particles may subsequently sink into the oscula of the sponges and block the exhalant flow, unless the oscula pump fast enough to prevent the particle from falling into the osculum. Third, large oscula may be advantageous for sponges living in stagnant water. At the shallow pristine site in this study, Meteor Bay, the absence of oceanic water currents leaves a mellow wave action as the major source of mixing and water movement. Sponges with few large oscula or with one single, giant osculum might break the stagnant oscillation by their strong oscular outflow, thus preventing re-ingestion of waste particles and promoting the influx of new particles. Analogously, it is known that an elongated osculum is a physiological adaptation of a sponge ensuring that the waste products are pushed away to a sufficient distance (Reiswig 1975).

If having large oscula would be an adaptation to living in stagnant water, one would expect to find large oscules as well in sponges living in deep stagnant waters, such as cave environments. We have so far not found oscula larger than 0.8 cm (diameter) in deep water (> 20 m) specimen of *C. reniformis*. It remains to be investigated to whether other factors, such as increased pressure, prevent sponges from making large oscula at greater depths.

### 4.3 Aquaculture perspectives

Looking from an aquaculture perspective, it becomes apparent that pumping rates of *C. reniformis* measured under pristine conditions are not a good proxy to calculate its direct bioremediation potential when these sponges are positioned in the vicinity of fed aquacultures such as open fish cages. On the other hand, there may be more food available in the water column near fish cages, which the sponges might eat at the same rate or even faster when compared to a pristine site. For example, if the pumping rate is three-fold lower and the food concentration is 6-fold higher, the sponges can potentially take up twice as much food as under pristine conditions and may thus grow faster. Indeed, Osinga et al. (2010) found that explants of *Dysidea avara* grew better under these polluted conditions than under pristine conditions. However, Osinga et al. (2010) also showed that *C. reniformis* is much more sensitive to high sediment levels than *D. avara. C. reniformis* explants cultured directly underneath the fish cages were completely smothered by fishfarm based organic matter, became soft and eventually died (Osinga et al. 2010). Notwithstanding this, we regularly encountered the presence of healthy *C. reniformis* colonies in the vicinity of fish farms during the current field observations. This suggests that the threshold value for particle density above which food particles completely inhibit pumping is being exceeded directly under the fish farms. Future studies should therefore compare growth rates and food retention efficiencies of *C. reniformis* under different levels of turbidity in order to assess the potential of this species as extractive component in IMTA. Comparative growth experiments on C. reniformis have recently been carried out at Gundogan Bay and Karaada Island by our team, which will be reported elsewhere.

Water depth is of high importance to aquaculture design, because depth apparently decreases the potential of *C. reniformis* to pump and, in the case of integrated culture, to take up the organic pollution caused by fed aquacultures. In this respect, it would be of interest to study whether the ability to make large oscula is a genetic or a fenotypical adaptation. If having large oscula is a genotypic trait, it is worth to select for those genotypes. However, if it is just fenotypic adaptation, the advantage might disappear when sponges are transplanted to other areas.

### 4.4 Concluding remarks

It can be concluded that video analysis of neutrally buoyant particles moving with the oscular outflow provides a simple, useful alternative to existing techniques to measure sponge pumping such as heated microthermistors (Pile 1996; Vogel 1974, 1977; Mendola 2008), ingenious balanced-level pumping chamber and laser-floating mirror projecting (Riisgard et al. 1993), particle tracking velocimetry (Schallpy et al 2007; Mendola 2008), the InEx technique (Yahel et. al 2003) and the fluorescent dye-video camera techniques (Savarese et. al 1997; Weiss et al. 2008). Using the new method, it was shown that for *C. reniformis*, oscular outflow had a location-dependent, in most cases positive relationship with oscular size. Turbidity and depth both affected sponge pumping in a negative way, but for the locations tested, the effect of depth was more profound than the effect of turbidity. Depth affected all parameters investigated except sponge size, whereas turbidity only affected specific pumping rates normalized to sponge surface area. In this study, we were not able to test potential interactions between effects of depth and turbidity on pumping, due to the absence of *C. reniformis* at greater depths in the polluted area. Hence, it remains to be studied how pumping activity of *C. reniformis* responds to different gradients and combinations of depth and turbidity. In addition, it is of interest to test if other sponge species show similar responses to depth and turbidity as *C. reniformis*.

## Acknowledgements

This study was financially supported by the European Commission, Grant Agreement no 266033 (project SPECIAL).

